# Protein Crowding and Cholesterol Increase Cell Membrane Viscosity in a Temperature Dependent Manner

**DOI:** 10.1101/2023.02.08.527594

**Authors:** Balázs Fábián, Ilpo Vattulainen, Matti Javanainen

## Abstract

Shear viscosity of lipid membranes dictates how fast lipids, proteins, and other membrane constituents travel along the membrane and rotate around their principal axis, thus governing the rates of diffusion-limited reactions taking place at membranes. In this framework, the heterogeneity of biomembranes indicates that cells could regulate these rates *via* varying local viscosities. Unfortunately, experiments to probe membrane viscosity at various conditions are tedious and error prone. Molecular dynamics simulations provide a luring alternative, especially now that recent theoretical developments enable the elimination of finite-size effects in simulations. Here, we use different equilibrium methods to extract the shear viscosities of lipid membranes from both coarse-grained and all-atom molecular dynamics simulations. We systematically probe the variables relevant for cellular membranes, namely membrane protein crowding, cholesterol concentration, and the length and saturation level of the lipid acyl chains, as well as temperature. Our results highlight that in their physiologically relevant ranges, cholesterol concentration, protein concentration, and temperature have significantly larger effects on membrane viscosity than lipid acyl chain length and unsaturation level. Our work also provides the largest collection of membrane viscosity values from simulation to date, which can be used by the community to predict the diffusion coefficients or their trends *via* the Saffman–Delbrück description. Additionally, diffusion coefficients extracted from simulations exploiting periodic boundary conditions must be corrected for the finite-size effects prior to comparison with experiment, for which the present collection of viscosity values can readily be used. Finally, our thorough comparison to experiments suggests that there is some room for improvement in the description of bilayer dynamics provided by the present force fields.

## 1 Introduction

In cell membranes, lateral and rotational diffusion coefficients, *D*^lat^ and *D*^rot^ respectively, are two central parameters to describe the rate at which a molecule samples its membrane environment due to thermal agitation. The larger the lateral (translational) diffusion coefficient, the larger membrane area the diffusing molecule can explore in unit time. Similarly, the larger the rotational diffusion coefficient, the faster the molecule in question randomly rotates around its principal molecular axis per unit time. In both cases, it is obvious that the viscosity of the membrane affects the rate of diffusion: in a syrup-like cell membrane of high viscosity, the diffusion will be dramatically slower than in membranes of low viscosity. In the case of colloids, this dependence manifests itself in such a way that the diffusion coefficient of a moving particle is inversely proportional to the viscosity of the substance surrounding it, in other words, the diffusion of the particle is regulated by the viscosity of the substance in its vicinity. Although cell membrane proteins are not actually colloids, it is reasonable to assume that a similar dependence, at least approximately, is also valid for integral membrane proteins, in which case the key question in terms of diffusion is how membrane viscosity behaves in complex biological conditions. The aim of this paper is to shed light on this question.

Knowing how the values of the lateral and rotational diffusion coefficients depend on the conditions in the host membrane—such as lipid composition, macromolecular crowding, or the interactions with the actin cytoskeleton—is crucial, because it helps to understand how proteins move to find one another, and how they rotate to find each other’s correct interaction interfaces to form functional protein–protein and protein–lipid units.^1–3^ Still, measuring either of these two diffusion coefficients is generally a tedious task.^4^ The experimental setups to measure *D*^lat^ often require the labeling of the proteins with fluorescent labels or nanoparticles^5^ that inevitably perturb the system.^6,7^ Several approaches exist to determine *D*^rot^, all with their own limitations.^8^ Notably, only a few techniques probe the short-time window required for the determination of *D*^rot^.^9,10^

Here, the celebrated Saffman–Delbrück (SD) model^11,12^ comes in handy, as it is in principle able to predict the values of both *D*^lat^ and *D*^rot^ from basic membrane properties, such as solvent and membrane shear viscosities (*μ*_f_ and *μ*_m_ respectively), membrane thickness (*h*), and the cross-sectional (in-plane) radius (*R*) of the membrane protein. The SD model predicts for lateral diffusion a relation

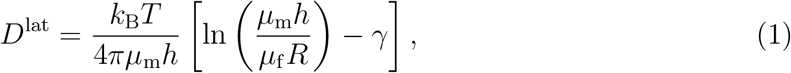

where *k*_B_ is the Boltzmann constant, *T* is temperature, and *γ* is the Euler–Mascheroni constant with a value of ≈0.577. The product of membrane shear viscosity and membrane thickness is often considered as a single parameter, the surface viscosity *η*, avoiding the requirement to know the membrane thickness since its precise and unique determination is not an easy task. For rotational diffusion, the SD model has the form

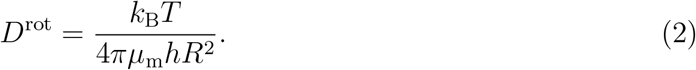

The SD model has been demonstrated to capture the logarithmic size-dependence of lateral diffusion in experiments^13–16^ as well as in computer simulations.^17–19^ Simulations have also confirmed the SD-like, quadratic size-dependence of *D*^rot^ in the case of rotational diffusion.^8,18,19^ Notably, the SD models of rotational and translational diffusion are both applicable for only small proteins whose radii are smaller than the SD length defined as *L*_SD_ = *hμ*_m_/(2*μ*_f_). In practice, this requirement means that *R/L*_SD_ < 0.1 must hold for the SD equations to be applicable.^20,21^ However, for large inclusions (or in the case of highly viscous surrounding fluid) the extensions of the SD model suggest scaling laws of *D*^lat^ ~ *R*^−1^ and *D*^rot^ ~ *R*^−3^, respectively,^11,12,21^ and similar relations have also been observed for membranes crowded with proteins using computer simulations.^8,17^

The matter that draws attention is that the SD formulas (Eqs. (1) and (2)) show how both lateral and rotational diffusion of membrane proteins are inversely proportional to membrane shear viscosity *μ*_m_, showing how the analogy to (Brownian) diffusion of colloids also applies qualitatively to the diffusion of membrane proteins. The fact that the values of these diffusion coefficients depend on membrane shear viscosity also provides a link to the control of biologically relevant operating conditions, since membrane viscosity in turn depends on temperature, membrane lipid composition, and the presence of other membrane-associated molecules such as membrane proteins. Unfortunately, experimental determination of membrane shear viscosity is challenging,^4^ and the literature values extracted with different experimental approaches are surprisingly scattered (see for example Appendix Table I in Ref. 22, which illustrates how different methods can produce membrane viscosity values that differ by a factor of about 200 for the same membrane composition at the same temperature). On the other hand, even if the differences in the values given by the different methods were paid less attention, the problem still remains that experimental studies in which the behavior of membrane viscosity as a function of temperature, lipid chain or head group composition, cholesterol concentration, or protein crowding has been systematically studied with some method are scarce,^22–25^ and the measured values are often complicated to interpret in the framework of the SD model.^26–28^

Considering the problems faced by experimental research, a natural alternative method for elucidating the behavior of membrane viscosity is biomolecular simulation. Computer simulations, such as classical molecular dynamics (MD) simulations, are often cheaper, less labor-intensive, and even faster to perform than lab experiments. Moreover, they can be implemented in circumstances where the composition of the systems and the thermodynamic conditions are unambiguously determined, so that the comparison with the predictions given by theories is as direct as possible. Thus, it is of significant interest to harness them in the determination of *D*^lat^, *D*^rot^, and *μ*_m_ under varying thermodynamic, biologically relevant conditions.

In this work, we extracted shear viscosity values for membranes at different lipid compositions, temperatures, and degrees of protein crowding using both coarse-grained (CG) and all-atom (AA) simulation models. The analysis of the simulation data was carried out rigorously, taking into account the finite size of the simulated systems and the effects produced by the periodic boundary conditions (PBCs) used in the simulations. For the CG systems, we first applied the SD models (Eqs. (1) and (2)) and their PBC-dependencies (Eqs. (3) and (4), see discussion below) to various membrane protein simulations, and extracted fairly consistent shear viscosity values in the limit of dilute membrane protein concentration. Next, with the most promising approaches that we identified based on analysis of lateral and rotational diffusion, we evaluated the effects of temperature and protein crowding on membrane shear viscosity. Finally, we probed lipid diffusion in all-atom simulations and used the PBC-dependence of lateral diffusion (Eq. (3)) to extract the viscosities of lipid bilayers that differ in the length and unsaturation level of the lipid chains, in cholesterol content, and in temperature.

The results show that the value of membrane shear viscosity can depend radically on the conditions prevailing in cell membranes. The effect of protein crowding is particularly important, as the membrane viscosity can easily increase by almost two orders of magnitude when the concentration of membrane proteins is increased from the dilute protein-poor limit to the protein-rich case. The corresponding effect, although weaker, is observed by increasing the concentration of cholesterol. Varying the unsaturation level and chain length of the lipid hydrocarbon chains has its own effect, but these effects are relatively weak. On the other hand, the effect of temperature on membrane viscosity is significant, which generally manifests itself as increasing the temperature decreases membrane viscosity. Biologically, these results are fascinating, especially regarding the effects of protein crowding and cholesterol, because cell membranes are known to be heterogeneous, meaning, among other things, that the local concentration of proteins and cholesterol varies significantly along the cell membrane. The viscosity of cell membranes is therefore never constant, but varies along the surface of the cell membrane depending on the local composition of the cell membrane.

Additionally, our results summarize the quality of different methods in extracting shear viscosities from membrane simulations, and provide a reference for the membrane viscosity values of numerous membranes that can be used to either estimate diffusion coefficients in such membranes, or to correct for PBC-induced effects.^29–33^ Unfortunately, our results also highlight the shortcomings of the Martini 2 and CHARMM36 force fields in quantitatively capturing the experimental values of diffusion coefficients and shear viscosities.

## 2 Methods

### 2.1 Accounting for Periodic Boundary Conditions

A key problem regarding simulations is related to the size of the systems being studied. While experimental data corresponds to the scale of cells in the range of tens of micrometers, in molecular simulations the size of the system is typically a few tens of nanometers. Given this, most modern biomolecular molecular dynamics (MD) simulations are performed using periodic boundary conditions (PBCs), which eliminate unwanted boundary effects of the 3D simulation box and ensure the conservation of total linear momentum. The use of PBCs gives the illusion that the simulated system is infinite, but the reality is that the small size of the system has drastic effects on the lateral and rotational dynamics of the simulated molecules, preventing straightforward comparison between simulation and experiment. Already in 2004, Yeh and Hummer derived a formula for the dependence of translational diffusion coefficient on simulation box size in isotropic systems,^34^ and a similar result for rotational diffusion was published recently.^35^ The main use of these formulas is to correct for the PBC-induced effects, and thus extract size-independent diffusion coefficients which can be directly compared to experiments. Recently, these concepts were extended to lipid membranes, as Vögele et al. derived models for both lateral^31,32^ and rotational^33^ diffusion of membrane-embedded objects such as proteins or lipids. Based on their work, for lateral diffusion in a flat simulation box with a membrane whose lateral dimension *L* (area of the simulated membrane being *L*^2^) is substantially larger than the thickness of the membrane *L_z_*, (*L* ≫ *L_z_*), and solvent layer thickness of 2*H*, the correction for the PBC-induced effects reads^31^

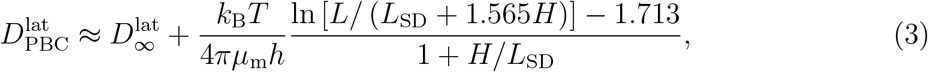

where 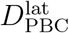 is the value in a simulation with PBCs and 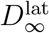 is the (corrected) value in an infinite system (referred to as *D*^0^ in some works^30^). While Eq. (3) is derived for membrane-spanning objects, it was also found to describe the diffusion of monotopic proteins and lipids to a great accuracy due to a strong coupling of the two leaflets.^32^ The corresponding formula for rotational diffusion in a membrane with an area of *A*_box_ is^33^

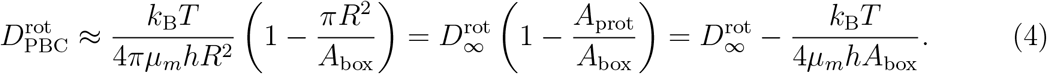

Here 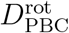 and 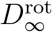 are again the values in the simulation with PBCs and in an infinite system, and *A*_prot_ is the cross-sectional area of the protein. These two elegant models were demonstrated to well capture the PBC-dependence of both *D*^lat^ and *D*^rot^.^31–33^ Eqs. (3) and (4) can also be extremely useful in calculating membrane viscosity *μ_m_* that is quite tedious to extract from MD simulations.^36,37^ Noteworthy, *μ_m_* is independent of system size, and therefore Einstein-like relations linking diffusion coefficients with membrane viscosity only hold in an infinite system. Under other conditions, corrections need to be made to these relations, such as Eq. (4) for *D*^rot^ (see also Refs. 29 and 30 for a PBC-corrected Saffman–Delbrück model for *D*^lat^).

Moving on, given that the membrane viscosity *μ_m_* is known, Eqs. (3) and (4) are useful in eliminating PBC-induced effects from diffusion coefficients extracted from simulations. In practice, this means that when the simulations measure the lateral and rotational diffusion coefficients in a finite-sized (small) system, Eqs. (3) and (4) can be used to determine the values that these diffusion coefficients would have in an infinite-sized system—which enables a reliable comparison of the simulation results with experimental data. Additionally, with the proper theoretical framework at hand, viscosity or the diffusion coefficients of the infinite system could be used as target parameters in force field development.

### 2.2 Coarse-Grained (CG) Simulations

For this work, we analyzed our simulation data described in our earlier work,^8,17^ as well as the data of new simulations performed for this work. The used CG simulations are listed in Table 1. The protein structures referred to in Table 1, their PDB identifiers, and the effective radii (*R*_eff_) used in fits of the SD models are shown in Fig. 1. The effective radii were determined in our previous work,^17^ and they describe the size of the diffusing entities, *i.e*. the proteins together with their stably-associated lipid shells. All CG simulations were performed in DPPC membranes, which remained in the fluid phase at all used simulation conditions.

**Table 1:** Table of simulated CG systems. “System” identifies the name of the system, “#Proteins” is the number of proteins in the simulation system, the lipid/protein ratio (describing crowding with proteins) is given per leaflet, and *t*_sim_ is the simulation time. The notation P[1–7] refers to all seven proteins. With temperature, 4 *T*s indicates that the simulation was repeated at four temperatures (300, 315, 330, and 345 K). In the case of 5 *T*s, an additional simulation at 360 K was performed. For systems labeled 1 T, only one simulation at 315 K was performed. “Data ref.” stands for the reference number that has the link to the deposited simulation data.

| System | # Proteins | Lipid/Protein ratio | $t_{\text{sim}}$ | Temperature | Data ref. |
| --- | --- | --- | --- | --- | --- |
| **Set CG-1: Dilute Polydisperse Membranes of Varying Size** |  |  |  |  |  |
| P-S | 1×P[1–7] | 400 | 80 $\mu\text{s}$ | 1 $T$ | 43 |
| P-M | 2×P[1–7] | 400 | 100 $\mu\text{s}$ | 1 $T$ | 44 |
| P-L | 8×P[1–7] | 400 | 60 $\mu\text{s}$ | 1 $T$ | 43 |
| **Set CG-2: Single-Protein Membranes at Various Temperatures** |  |  |  |  |  |
| S-1 | 1×P1 | 400 | 20 $\mu\text{s}$ | 4 $T$s | 45,46 |
| S-2 | 1×P2 | 400 | 20 $\mu\text{s}$ | 4 $T$s | 45,46 |
| S-3* | 1×P3 | 400 | 20 $\mu\text{s}$ | 4 $T$s | 45,46 |
| S-4** | 1×P4 | 400 | 20 $\mu\text{s}$ | 4 $T$s | 45,46 |
| S-5 | 1×P5 | 400 | 20 $\mu\text{s}$ | 4 $T$s | 45,46 |
| S-6 | 1×P6 | 400 | 20 $\mu\text{s}$ | 4 $T$s | 45,46 |
| S-7 | 1×P7 | 400 | 20 $\mu\text{s}$ | 4 $T$s | 45,46 |
| **Set CG-3: Single-Protein Membranes at Various Protein/Lipid Ratio** |  |  |  |  |  |
| S-3<sup>50</sup> | 1×P3 | 50 | 50 $\mu\text{s}$ | 1 $T$ | 47 |
| S-3<sup>75</sup> | 1×P3 | 75 | 50 $\mu\text{s}$ | 1 $T$ | 47 |
| S-3<sup>100</sup> | 1×P3 | 100 | 50 $\mu\text{s}$ | 1 $T$ | 47 |
| S-3<sup>200</sup> | 1×P3 | 200 | 50 $\mu\text{s}$ | 1 $T$ | 47 |
| S-3<sup>400*</sup> | 1×P3 | 400 | 50 $\mu\text{s}$ | 1 $T$ | 45,47 |
| **Set CG-4: Single-Protein Membranes at Various Temperatures and Sizes** |  |  |  |  |  |
| S-4<sup>400**</sup> | 1×P4 | 400 | 20 $\mu\text{s}$ | 5 $T$s | 45,46 |
| S-4<sup>1600</sup> | 1×P4 | 1600 | 20 $\mu\text{s}$ | 5 $T$s | 48 |
| S-4<sup>3600</sup> | 1×P4 | 3600 | 20 $\mu\text{s}$ | 5 $T$s | 49 |
| **Set CG-5: Crowded Monodisperse Membranes at Various Temperatures and Sizes** |  |  |  |  |  |
| M-4<sup>200</sup> | 4×P4 | 50 | 20 $\mu\text{s}$ | 5 $T$s | 50 |
| M-4<sup>450</sup> | 9×P4 | 50 | 20 $\mu\text{s}$ | 5 $T$s | 51 |
| M-4<sup>1800</sup> | 36×P4 | 50 | 20 $\mu\text{s}$ | 5 $T$s | 52 |
| */** Same system |  | total | 1.45 ms |  |  |

**Figure 1:**
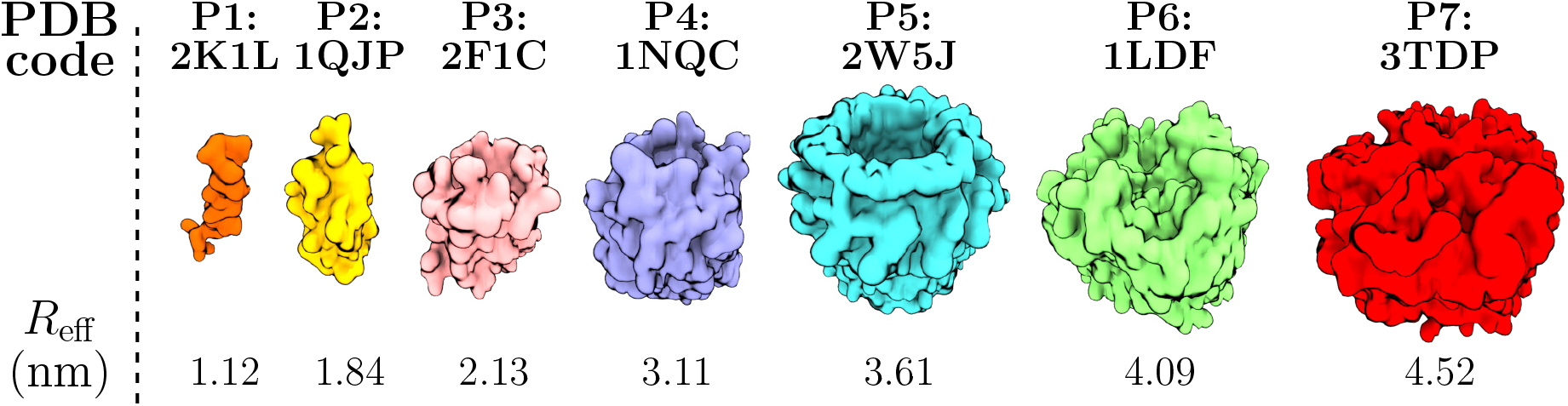
Structures, PDB identifiers, and effective radii of the seven proteins (P1–P7) explored in this work through CG simulations.

#### Set CG-1: Dilute Polydisperse of Varying Size

We analyzed our previously generated simulation data of a dilute membrane (400 lipids per protein per membrane leaflet) with seven protein types of varying radii “P-M” (standing for Polydisperse set of proteins, Medium size) in Table 1.^8,17^ The used proteins are shown in Fig. 1 with further details available in Ref. 17. The membrane contained two copies of each of the protein types and consisted of ~300,000 coarse-grained beads with a membrane edge length of ~61 nm. Additionally, for this work, we also simulated a 4 times Larger (“P-L”, box edge ~ 122 nm) and 50% Smaller (“P-S”, box edge ~43 nm) versions of these membranes that contained 8 or 1 copies of each of the protein types, respectively.

#### Set CG-2: Single-Protein Membranes at Various Temperatures

We also considered dilute single-protein systems for each of these seven proteins. These systems are numbered based on the seven proteins (1–7 with increasing size) shown in Fig. 1. The single-protein simulations contained 400 lipids per protein per leaflet, and were performed at 300, 315, 330, and 345 K. They are labeled as “S-” (standing for Single) in Table 1. The single-protein simulation systems are described in Ref. 17, and were extended here to different temperatures.

#### Set CG-3: Single-Protein Membranes With Varying Levels of Protein Crowding Described in Terms of the Protein/Lipid Ratio

Additionally, the simulation in Set CG-2 described above at 315 K was repeated for protein number 3 with varying lipid-to-protein ratios (50, 75, 100, 200, and the original 400). These systems are labeled “S-3^no.lipids^” in Table 1. The single-protein simulation systems are described in Ref. 17, and were extended here to different lipid amounts.

#### Set CG-4: Single-Protein Membranes At Various Temperatures and Sizes

For protein number 4 in Set CG-2, the simulations at 300, 315, 330, 345, and 360 K were also repeated with 4 and 9 times larger membranes (yet always with a single protein), containing 1600 and 3600 lipids per protein per leaflet, respectively. These systems are labeled “S-4^no.lipids^” in Table 1. Again, the single-protein simulation systems are described in Ref. 17, and were extended here to different temperatures and system sizes.

#### Set CG-5: Crowded Monodisperse Membranes At Various Temperatures and Sizes

Finally, we simulated crowded (50 lipids per protein per leaflet) membranes with protein number 4 in three sizes. The membranes contained 4 (edge length ~15 nm, 400 lipids), 9 (edge length ~22 nm, 900 lipids), or 36 (edge length ~44 nm, 3600 lipids) proteins, and the simulations were performed at 300, 315, 330, 345 K, and 360 K. These simulations are labeled “M-4^no.lipids^” in Table 1. The M-4^900^ simulation system at 315 K is described in Ref. 17, and were extended here to different system sizes and temperatures.

For a complete list of CG simulations with their duration, see Table 1. In all analyses, the first 1 *μ*s of the simulations was discarded. We used the MARTINI 2.2 force field^38–40^ with reduced Lennard-Jones interactions to prevent excessive protein aggregation.^8,17,41^ The CG simulations were performed with GROMACS versions 5.0.x (original simulations in Ref. 17) and 2018.4 (newer simulations).^42^ The simulation methods, thoroughly explained in Ref. 17, were applied also for all new simulations. The references to simulation data, available online, are also provided in the “Data ref.” column in Table 1, and these uploads include simulation parameter files (mdp).

### 2.3 All-Atom (AA) Simulations

We also performed simulations of protein-free lipid bilayers in all-atom (atomistic) resolution. In these sets we varied either the lipid acyl chain length and/or saturation, temperature, or cholesterol content. They are all listed in Table 2, and classified below.

**Table 2:** Table of simulated all-atom systems. The system name, the numbers of lipids in small/medium/large systems, and simulation temperature are given. The membrane viscosities extracted from the the system-size dependence of lateral diffusion (Eq. (3)) are given. The references to the uploaded simulation data are also provided.

| System | # Lipids | $T$ (K) | $\mu_m$ (mPa·s) | Data ref. |
| --- | --- | --- | --- | --- |
| **Set AA-1: Varying Acyl Chains** |  |  |  |  |
| DMPC | [64/256/1024] DMPC | 333 | 11.5 | 57 |
| DRPC | [64/256/1024] DRPC | 333 | 9.8 | 57 |
| DPPC | [64/256/1024] DPPC | 333 | 15.1 | 57 |
| PYPC | [64/256/1024] PYPC | 333 | 12.8 | 57 |
| DYPC | [64/256/1024] DYPC | 333 | 11.1 | 57 |
| DSPC | [64/256/1024] DSPC | 333 | – | 57 |
| SOPC | [64/256/1024] SOPC | 333 | 15.6 | 57 |
| SLiPC | [64/256/1024] SLiPC | 333 | 13.7 | 57 |
| DOPC* | [64/256/1024] DOPC | 333 | 16.8 | 57 |
| DLiPC | [64/256/1024] DLiPC | 333 | 11.0 | 57 |
| **Set AA-2: DOPC with Varying Cholesterol Concentrations** |  |  |  |  |
| DOPC<sup>0%</sup>* | [64/256/1024] DOPC<br>+ [0/0/0] CHOL | 333 | 16.8 | 57 |
| DOPC<sup>11%</sup> | [64/256/1024] DOPC<br>+ [8/32/128] CHOL | 333 | 17.0 | 58 |
| DOPC<sup>20%</sup> | [64/256/1024] DOPC<br>+ [16/64/256] CHOL | 333 | 15.6 | 58 |
| DOPC<sup>29%</sup> | [64/256/1024] DOPC<br>+ [26/104/416] CHOL | 333 | 18.0 | 58 |
| DOPC<sup>38%</sup> | [64/256/1024] DOPC<br>+ [40/160/640] CHOL | 333 | 22.0 | 58 |
| DOPC<sup>47%</sup> | [64/256/1024] DOPC<br>+ [56/224/896] CHOL | 333 | 38.7 | 58 |
| **Set AA-3: POPC with Varying Cholesterol Concentrations** |  |  |  |  |
| POPC<sup>0%</sup> | [64/256/1024] POPC<br>+ [0/0/0] CHOL | 298 | 50.7 | 59 |
| POPC<sup>11%</sup> | [64/256/1024] POPC<br>+ [8/32/128] CHOL | 298 | 54.4 | 59 |
| POPC<sup>20%</sup> | [64/256/1024] POPC<br>+ [16/64/256] CHOL | 298 | 71.6 | 59 |
| POPC<sup>29%</sup> | [64/256/1024] POPC<br>+ [26/104/416] CHOL | 298 | 83.3 | 59 |
| POPC<sup>38%</sup> | [64/256/1024] POPC<br>+ [40/160/640] CHOL | 298 | 129.3 | 59 |
| POPC<sup>47%</sup> | [64/256/1024] POPC<br>+ [56/224/896] CHOL | 298 | 255.4 | 59 |
| **Set AA-4: DOPC at Various Temperatures** |  |  |  |  |
| DOPC<sup>293 K</sup> | [64/256/1024] DOPC | 293 | 65.4 | 60 |
| DOPC<sup>303 K</sup> | [64/256/1024] DOPC | 303 | 38.4 | 60 |
| DOPC<sup>313 K</sup> | [64/256/1024] DOPC | 313 | 28.0 | 60 |
| DOPC<sup>323 K</sup> | [64/256/1024] DOPC | 323 | 16.7 | 60 |
| DOPC<sup>333 K*</sup> | [64/256/1024] DOPC | 333 | 16.8 | 57 |

\* Same system
total 75 μs

#### Set AA-1: Varying Acyl Chains

We set up bilayers composed of phosphatidylcholine (PC) lipids with varying acyl chains. The original bilayers had a total of 64 lipids. By replicating their coordinates in the membrane plane, bilayers with 256 and 1024 lipids were also created. The considered lipids and their acyl chains were DMPC (14:0,14:0), DRPC (14:1,14:1), DPPC (16:0,16:0), PYPC (16:0,16:1), DYPC (16:1,16:1), DSPC (18:0,18:0), SOPC (18:0,18:1), SLiPC (18:0,18:2), DOPC (18:1,18:1), and DLiPC (18:2,18:2). Here, we mainly use the CHARMM-GUI nomenclature, but use consistently “Li” for linoleyl, whereas its naming varies in CHARMM-GUI. These membranes were simulated at 333 K to ensure that all of them were in the liquid-disordered phase. These systems are labeled with the respective lipid name in Table 2.

#### Set AA-2: DOPC with Varying Cholesterol Concentrations

Additionally, we performed simulations on mixtures of DOPC and cholesterol, with cholesterol molecules included in addition to the 64/256/1024 DOPC molecules. The cholesterol concentrations were 11 mol% (8 molecules in the smallest system), 20 mol% (16 molecules), 29 mol% (26 molecules), 38 mol% (40 molecules), and 47 mol% (56 molecules). These membranes were simulated at 333 K to be consistent with Set AA-1. These systems are labeled as DOPC^cholesterol concentration^ in Table 2.

#### Set AA-3: POPC with Varying Cholesterol Concentrations

In addition to DOPC in Set AA-2, we also considered mixtures of POPC with cholesterol. The same amounts of cholesterol were used, but the simulations were performed at 298 K. These systems are labeled as POPC^cholesterol concentration^ in Table 2.

#### Set AA-4: DOPC Simulations at Various Temperatures

Finally, we repeated the DOPC simulation in Set AA-1 (at 333 K) at various other temperatures, namely at 293, 303, 313, and 323 K, all of which are above the main transition temperature of DOPC. These systems are labeled as DOPC^temperature^ in Table 2.

All membranes were solvated by 50 water molecules per lipid, and all simulations were 1 μs long with the first 10 ns excluded from the analyses. All in all, we performed 75 μs of atomistic simulations. For all-atom simulations, GROMACS 2020.x^42^ was used. The CHARMM36 force field^53,54^ was used together with the CHARMM-specific TIP3P model.^55,56^ For details on the simulation parameters, see SI. The simulation data, along with simulation parameter files (mpd) are provided online, and the references are provided in Table 2.

### 2.4 Analysis Methods

The 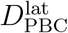 and 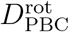 were extracted from our simulations affected by PBCs as in our earlier works.^8,17^ Briefly, we fitted

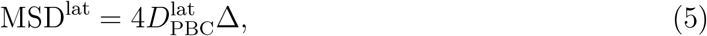

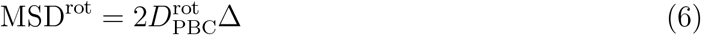

to lateral and rotational mean-squared deviation (MSD) data calculated as a function of lag time Δ, to reach the regimes of normal diffusion. The units of MSD data are length squared (lateral diffusion) and radians squared (rotational diffusion), hence the units of diffusion coefficients are cm^2^/s (lateral diffusion) and rad^2^/s = 1/s (rotational diffusion). For CG simulations, MSD was averaged over time and proteins of the same type (where applicable). For lateral diffusion, diffusion coefficients were extracted from fits to the lag time interval of 20–40 ns. For rotational diffusion, we used a lag time interval of 0.1–1 *μ*s for the “P” systems (Set CG-1), whereas for single-protein systems (Sets CG-2 and CG-3) we used a fitting interval of of 10–100 ns. Because only DPPC membranes were used in all the CG simulations, we adopted a single membrane thickness value of 4 nm. Viscosity of CG water as a function of temperature was extracted using the transverse current autocorrelation approach as implemented in `gmx tcaf`. The fitting intervals for the extraction of diffusion coefficients in the CG systems are listed in Table 3.

**Table 3:** Membrane shear viscosities of the coarse-grained DPPC bilayer (in mPa·s) at 315 K extracted using different approaches. PBC corrections are included, when they were found to be significant.

| **System** | **Data & Fit** | **$\mu_m$** |
| --- | --- | --- |
| **SD fit to $D_{\text{PBC}}^{\text{rot}}(R)$ vs. $R$, Eq. (2)** |  |  |
| P-S (Set CG-1) | Fig. S5 | 6.6 |
| P-M (Set CG-1) | Fig. S5 | 4.4 |
| P-L (Set CG-1) | Fig. S5 | 5.5 |
| **SD fits to $D_{\infty}^{\text{lat}}(R)$ & $D_{\infty}^{\text{rot}}(R)$ vs. $R$, Eqs. (3) & (4), and Eqs. (1) & (2)** |  |  |
| $D_{\infty}^{\text{rot}}$, S- (Set CG-2) | Fig. S7 | 4.7 |
| $D_{\infty}^{\text{lat}}$, P- (Set CG-1) | Fig. S4 | 6.9 |
| $D_{\infty}^{\text{lat}}$, S- (Set CG-2) | Fig. S6 | 4.6 |
| **Fit to $D_{\text{PBC}}^{\text{lat}}(L)$ vs. $L$, Eq. (3)** |  |  |
| P- (Set CG-1) | Fig. S1 | 11.3 |
| S-4 (Set CG-4) | Fig. S3 | 7.5<sup>†</sup> |
| **Fit to $D_{\text{PBC}}^{\text{rot}}(L)$ vs. $L$, Eq. (4)** |  |  |
| S-3 (Set CG-3) | Fig. S2 | 13.3 |

<sup>†</sup> At 315 K; more values at different temperatures shown in Fig. 2.

For the all-atom simulations, we fitted the leaflet-wise time- and ensemble-averaged MSD curves in the lag time interval between 10 and 100 ns. This region has sufficient statistics, while also representing normal diffusion. The mean and difference of the values calculated for the two leaflets were reported as the value and the error estimate, respectively. For the all-atom simulations, the membrane thicknesses were extracted from Gaussian fits to phosphorus density profiles. These thicknesses as well as box sizes normal to the membrane (and hence the thickness of the water slab *H*) were averaged over the three system sizes for each composition/temperature. The viscosities of the CHARMM-specific TIP3P water (also known as TIPS3P) at different temperatures were taken from Ref. 61. We also extracted the diffusion coefficients of POPC in POPC/cholesterol mixtures using the corrected unwrapping scheme of *NPT* trajectories^62^ and the generalized least squares (GLS) approach^63^ to ensure that the heuristic unwrapping scheme used by GROMACS and the fixed fitting interval of the MSD curves do not lead to major systematic errors. Indeed, the diffusion coefficient values extrapolated to the infinite system size using Eq. (3) differ by 4.3±5.7% between the heuristic default unwrapping scheme and the GLS tool, and the unwrapping error will be even smaller in the CG simulations with larger lateral dimensions. Thus, we extracted all diffusion coefficients in a conventional way.

## 3 Results and Discussion

### 3.1 Different Approaches Provide Similar Membrane Viscosities

Based on the above presentation, it is clear that there are several ways to determine the membrane viscosity, some of which are based on the examination of lateral diffusion and its PBC corrections, some on rotational diffusion including its PBC corrections, and in both cases there are several variations to implement the data analysis. The first goal is thus to ensure the functionality of the methodology and that the results given by the different methods are in line with each other with sufficient accuracy. Biologically relevant results regarding, for example, the crowding effects and the effect of cholesterol are discussed separately in the sections below.

We extracted membrane viscosities from single- and multi-protein CG simulations with the use of the SD models (Eqs. (1) and (2)) and their PBC corrections (Eqs. (3) and (4)). These results are reported in Table 3.

First, for every system we explored, we analyzed the diffusion coefficients 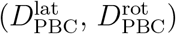 from the trajectories of the simulations using Eqs. (5) and (6). Next, we used the SD models (Eqs. (1) and (2)) to fit the 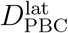 and 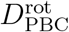 values as a function of protein size in systems where this approach was suitable.

We applied the SD model for lateral diffusion to the “P” membranes of Set CG-1, where the SD model was fitted to the lateral diffusion coefficients of the seven differently-sized proteins. The fits were performed separately for all three sizes of the simulation system (S, M, L). Data and fits are shown in Fig. S4. Not surprisingly, except in the limit of large proteins, the fit of the SD model to the lateral diffusion data is not very good if the data have not been corrected for PBC-induced effects, and thus the extracted viscosities are omitted from Table 3. Next, we applied the SD model for rotational diffusion to the same data (Set CG-1) on dilute membranes with a polydisperse set of proteins (data and fits shown in Fig. S5). The different system sizes provide fairly consistent values for membrane viscosity between 4.4 and 6.6 mPa s (see the top segment of Table 3), indicating that the PBC effects were insignificant for rotational diffusion at this system size regime, in line with Ref. 33.

Moving on to single-protein systems (“S-”, Set CG-2), we again applied the SD models to lateral and rotational diffusion coefficients as a function of protein size, but now also to data obtained at various temperatures. Here, both lateral and rotational diffusion coefficients were PBC-corrected using Eqs. (3) and (4), using the membrane viscosity extracted from system-size dependence of single-protein systems (discussed in the next section) and a geometric correction, respectively. Despite this correction, the lateral diffusion values are somewhat poorly fitted at higher temperatures (data and fits shown in Fig. S6), yet the value extracted at 315 K, 4.6 mPa s, agrees well with that from rotational diffusion, 4.7 mPa s.

Next, we fitted the PBC corrections of lateral and rotational diffusion to the 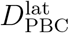 and 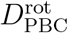 values extracted from simulations of different system sizes. Equation (3) describing the size-dependence of lateral diffusion was applied to the membranes with a polydisperse “P-” set of proteins simulated in three sizes (Set CG-1, data shown in the top panel of Fig. S1), which provided a viscosity value of 11.3 mPa s. As shown in the bottom panel of Fig. S1, the size-dependence of rotational diffusion is insignificant at large system sizes. Thus, we fitted the box-size dependence of 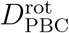 (Eq. (3)), to “S-3” systems (Set CG-3, data and fit in Fig. S2). The fit quality was subpar, and provided a somewhat higher value of 13.3 mPa s. The PBC corrections of 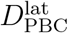 were also applied to the single-protein “S-4” systems simulated in three sizes (Set CG-4, data and fits shown in Fig. S3). This provided a value of 7.5 mPa s at 315 K (see the third and fourth segments of Table 3).

Finally, the fits of PBC correction, Eq. (3), to the 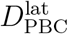 values extracted from “P-” systems (Set CG-1) (Fig. S1) provide the corrected values 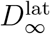. These values were fitted with the SD model, Eq. (1). This fit, shown in Fig. S4 along with the data, provides a viscosity value of 6.9 mPa s (see the second segment of Table 3).

All in all, the spread of the obtained values (7.2±3.1 mPa s) is remarkably small considering the uncertainties in all fitting parameters. Moreover, while all studied systems consist of DPPC lipids with a dilute concentration of proteins, they still differ in the nature of these proteins; Set CG-1 has 7 kinds of proteins present, each system in Set CG-2 contains a different protein, whereas Sets CG-3 and CG-4 have only one (but different) protein present. Thus, it seems that despite the probe (type(s) of embedded protein(s)), the used approaches provide consistent results for the viscosity of the underlying lipid matrix. Additionally, as discussed in the introduction, the values extracted using traditional SD models are affected by PBC effects. Still, these effects seem to be relatively small in our P systems as they fortuitously fall close to the zero error contour of PBC effects.^30^ The extracted shear viscosity values (7.2±3.1 mPa s) agree well with earlier estimates of 5±0.1 mPa s obtained for the same CG DPPC model at a slightly higher temperature of 323 K using non-equilibrium simulations.^37^ We conclude that the analysis methods used here are consistent with sufficient accuracy.

### 3.2 Crowding Increases Membrane Viscosity in a Temperature-Dependent Manner

Next, we extracted the temperature-dependence of membrane viscosity experienced by lateral and rotational motion. To this end, we considered the two approaches from the previous section that we believe to provide the most reliable data; we discarded fits of the SD models to lateral diffusion data across multiple protein sizes (*D*^lat^(*R*) vs. *R*, Eq. (1)), as finite size effects are hard to eliminate consistently (See Fig. S4). In the same manner, the finite size correction for rotational diffusion (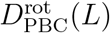 vs. *L*, Eq. (4)) suffers from poor statistics (See Fig. S2). The quality of the fits of the size-dependence of rotational diffusion (*D*^rot^(*R*) vs. *R*, Eq. (2)) to the values measured from polydisperse “P-” set of proteins (Set CG-1) was also subpar (See Fig. S5).

To overcome these limitations, we first considered the 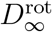 values measured at different temperatures (Set CG-2) in the single-protein (dilute concentration) case and shown in Fig. S7. For rotational diffusion, correcting for finite size effects is simple as it purely depends on the protein and membrane areas (Eq. (4)). Thus, we fitted the SD model, Eq. (2) to these PBC-corrected data (Eq. (4)), and extracted *μ*_m_ as a function of temperature from these fits. These data are shown in Fig. 2 in red.

**Figure 2:**
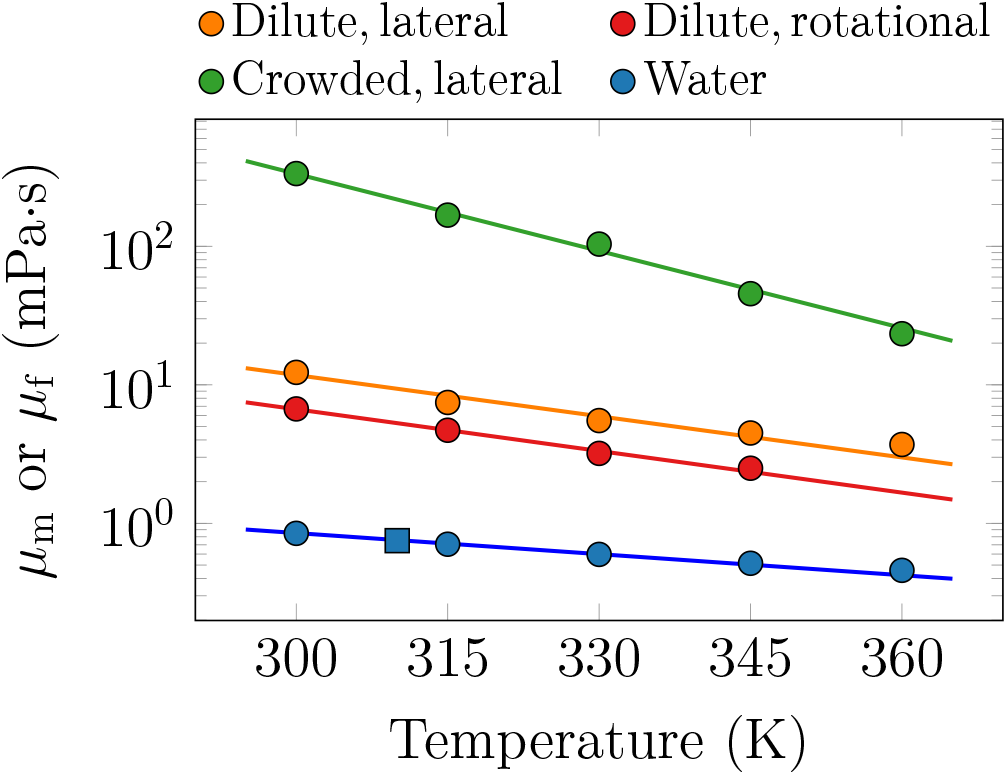
DPPC membrane (*μ*_m_) and solvent (*μ*_f_) viscosities as a function of temperature based on the CG simulation models. Values for water are calculated from transverse current autocorrelation (circles) or from the Green–Kubo equation (square) and shown in blue. The membrane viscosity is obtained in two ways: a) the values based on fits of the SD equation (Eq. (2)) to the PBC-corrected data (Eq. (4)) on rotational diffusion coefficients of proteins in dilute single-protein systems (Set CG-2) are shown in red; b) the values arising from fits of the size dependence of lateral diffusion of proteins (Eq. (3)) in dilute single-protein systems (set CG-4) are shown in orange. Finally, the values based on similar fits of the size-dependence of lateral diffusion of proteins but in crowded multi-protein systems (Set CG-5) are shown in green. Note the logarithmic scale on the ordinate.

For temperature dependence of viscosity experienced by lateral diffusion in the dilute protein-poor case, we extended the analysis based on PBC correction for lateral diffusion to temperatures other than 315 K (“S-4” systems, Set CG-4). The data and fits of Eq. (3) at each temperature are shown in the top panel of Fig. S3). In the fits, we used the values of *μ*_f_ of Martini water that we calculated using the transverse current autocorrelation approach.^64^ We also checked that the Green–Kubo-based method provided similar values for *μ*_f_. The calculated values of *μ*_f_ and *μ*_m_ are shown in Fig. 2 in orange and blue, respectively.

We also performed similar fits of Eq. (3) to the data extracted from membranes crowded with protein P4 (“M-4 systems”, Set CG-5). The data and fits are shown in the bottom panel of Fig. S3. Notably, this demonstrates the power of Eq. (3) in extracting the viscosities of crowded membranes, as the SD model breaks down in crowded systems.^8,17^

As suggested by Fig. 2, *μ*_m_ has an exponential temperature dependence in both dilute and crowded membranes. The chosen approaches to extract *μ*_m_ from lateral and rotational diffusion provide slightly different values of *μ*_m_, which is not surprising considering the subpar quality of the fits to rotational diffusion data in Fig. S7. Still, the significantly larger disagreements between the viscosities observed by the two types of motion in experiments in cellular membranes likely stem from some specific interactions of the probed molecule with, *e.g*., the actin cytoskeleton.^26^

The viscosity of the crowded membranes (Set CG-5, 50 lipids per protein per leaflet) is ≈27-fold higher than the viscosity of dilute protein-poor membranes at 300 K, yet the ratio decreases to ≈6 at 360 K. These different temperature dependencies suggest that membrane viscosity in the crowded membrane consists of two components: the temperature-dependent viscosity of the lipids, and a temperature-independent geometric exclusion effect.

The conclusion can be drawn from the results that protein crowding plays an exceptionally significant role in the viscosity of cell membranes. Particularly high viscosity is evident in protein-rich membrane regions where the local concentration of proteins is high, while protein-poor regions where the concentration of proteins is low are significantly less viscous.

### 3.3 Lateral and Rotational Diffusion Have Similar Temperature Dependence

Next, we evaluated whether the temperature dependencies of lateral and rotational diffusion are similar, and whether they depend on the protein size. We used the PBC-corrected 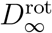 and 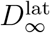 values for the seven single-protein systems (Set CG-2), with the seven proteins shown in Fig. 1. The 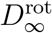 values were also used in the determination of the viscosity in Fig. 2. For the correction in 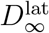, we used the membrane viscosity values extracted using Eq. (3) on Set CG-4 (orange curve in Fig. 2). The natural logarithm of the resulting 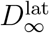 and 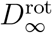 values are shown in Fig. 3 as a function of inverse temperature, which allows for the extraction of the activation energy *E*_A_ by fitting the data to the Arrhenius equation

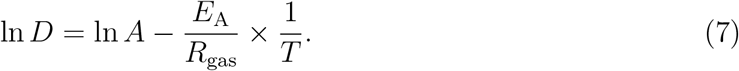

Here, *R*_gas_ the universal gas constant (not to be confused with the radius *R*), and *A* a temperature-independent pre-factor. The diffusion coefficients had units of cm^2^/s and rad^2^/s (=1/s), yet this only affects the vertical positioning of the curves and not the extracted activation energies.

**Figure 3:**
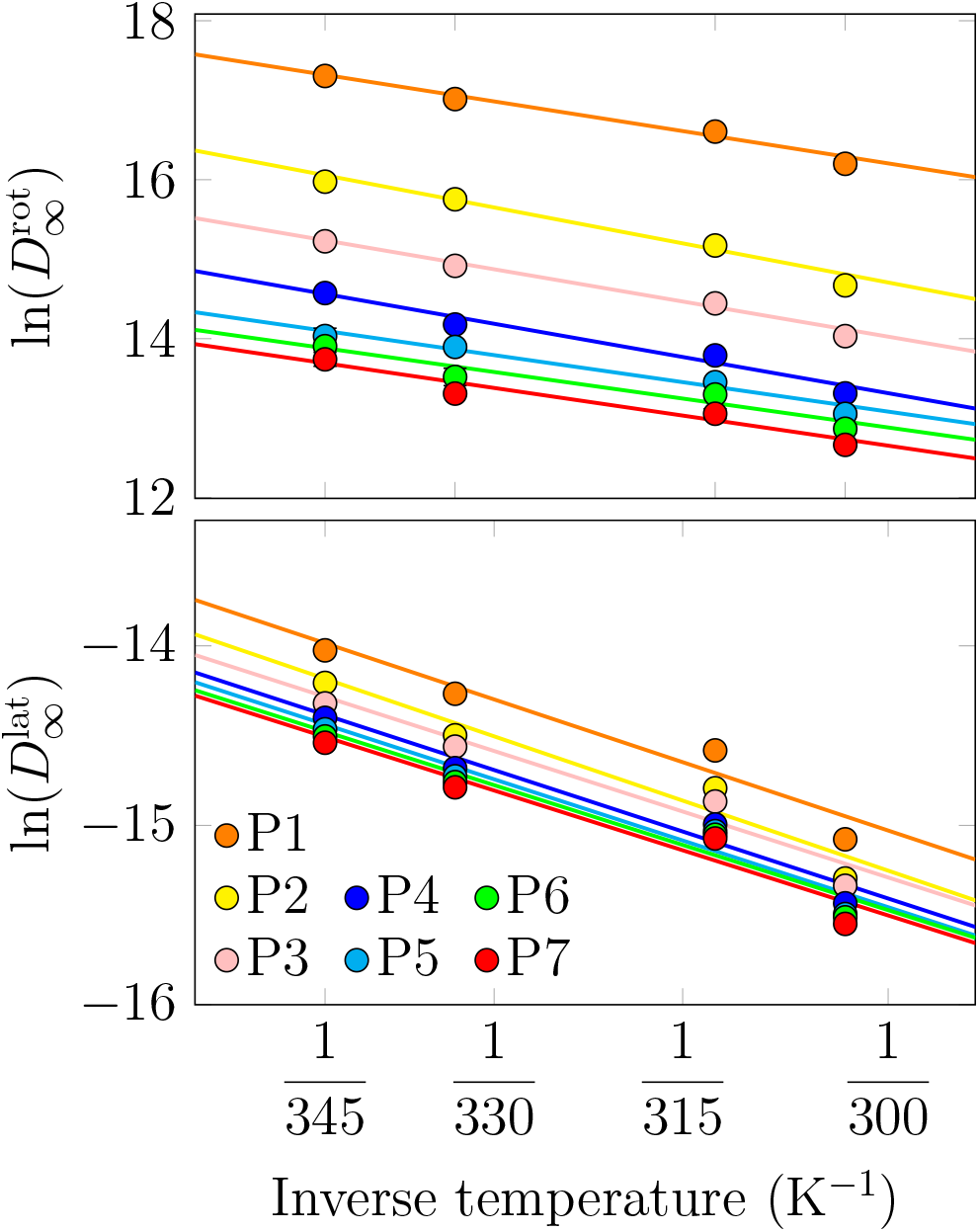
Rotational (top) and lateral (bottom) diffusion coefficients of the membrane proteins P1–P7 in the CG single-protein systems (“S” in Table 1) as a function of simulation temperature. The data are plotted in the Arrhenius fashion. Straight lines are fits of Eq. 7 to systems in the fluid phase. The rotational diffusion coefficients are PBC-corrected based on the protein and membrane areas according to Eq. (4), whereas the lateral diffusion coefficients were corrected using Eq. (3) and the membrane and solvent viscosity values in Fig. 2.

The Arrhenius form can be well fitted to the rotational diffusion coefficient values (Fig. 3), and the extracted activation energies are provided in Table 4. Curiously, these values show no clear dependence on the protein size. In terms of activation energies of the viscosities experienced by proteins in experiments, Bigelow et al. measured a value of *E*_a_ ≈ 47 kJ/mol for Ca-ATPase in sarcoplasmic reticulum membranes.^65^ This value is about twice the value observed in our simulations, which could be due to numerous reasons, such as the fact that the cell membrane used in experiment is much more complex than the simplified model membrane used in the simulation model.

**Table 4:**
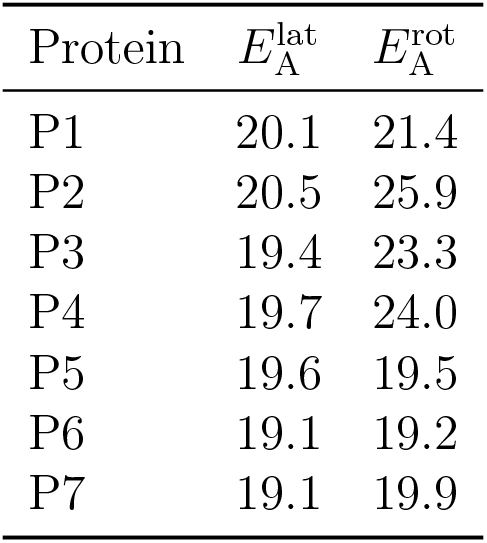
Activation energies (in kJ/mol) for lateral and rotational diffusion extracted from CG simulation data in Fig. 3 using Eq. (7).

We also extracted lateral diffusion coefficients of the proteins from the same systems. The data for the fluid phases were again well fitted by the Arrhenius form, and the activation energies did not display any clear dependence on the protein size. What is more, the activation energies for lateral motion were very similar to those extracted using data for rotational diffusion. Experimentally, the activation energy for lateral diffusion of rhodopsin in DMPC membranes was measured to be 46.9 kJ/mol, whereas a value of 16.2 kJ/mol was measured for Ca-ATPase in sarcoplasmic reticulum membranes, and values of 33.5 kJ/mol and 25.0 kJ/mol were obtained for acetylcholine receptor in DMPC and soybean membranes, respectively.^66^ These values are in reasonable agreement with our simulations, considering the differencies in membrane compositions, and the limitations of the CG models in describing dynamics and temperature-dependencies of membrane properties.^67^

The clear exponential dependencies of both rotational and lateral diffusion coefficients on temperature is on the one hand expected but on the other hand surprising. It is expected in the sense that the Arrhenius form can be fitted to practically any data as long as the temperature range is narrow enough, and then the activation energy describes how quickly the diffusion coefficient increases in this temperature range as the temperature rises. On the other hand, the observed exponential dependencies are somewhat surprising given that the SD formalisms for rotational and lateral diffusion, Eqs. (2) and (1), have explicit linear dependencies on temperature. This indicates that the implicit exponential dependence of membrane viscosity on temperature (Fig. 2) dominates over the explicit temperature-dependencies in the SD equations at temperatures that are of interest for biological applications. Indeed, we extracted the activation energies from the viscosity values shown in Fig. 2 and obtained similar values of 17.6 kJ/mol for the lateral motion and 19.2 kJ/mol for rotational motion. A recent fluorescence study found a value of ≈26.3 kJ/mol for the activation energy of the viscosity of large unilamellar vesicles (LUVs) composed solely of DPPC,^68^ which lies in the same ballpark as some of the experimental estimates for the activation energy of lateral diffusion listed above. Crowding, on the other hand, had a significant influence on the activation energy, and a fit to the data in Fig. 2 provided a value of 39.6 kJ/mol, similar to the higher estimates extracted for lateral diffusion using experiments.^66^

Based on these data, it is obvious that the membrane viscosity sensed by both lateral and rotational diffusion is the same, or similar.

### 3.4 Atomistic Simulations Capture Membrane Viscosity and Its Temperature Dependence

Next, we moved on to analyze our atomistic simulation data. The lateral diffusion coefficients extracted for phospholipids in the all-atom systems of increasing size are shown in Fig. S8 in the SI. Data for DSPC (18:0,18:0) has been omitted, as these 256- and 1024-lipid bilayers underwent a phase transition to a gel phase during the simulations.

The membrane shear viscosity values for DOPC as a function of temperature were extracted from Set AA-4 using Eq. (3). The resulting values are shown in Fig. 4, and the corresponding diffusion coefficients, as well as the ones extrapolated to infinite systems, and the ones obtained from pulsed field gradient NMR experiment,^69,70^ are all shown in Fig. S10 as a function of temperature. The viscosity values are provided in Table 2.

**Figure 4:**
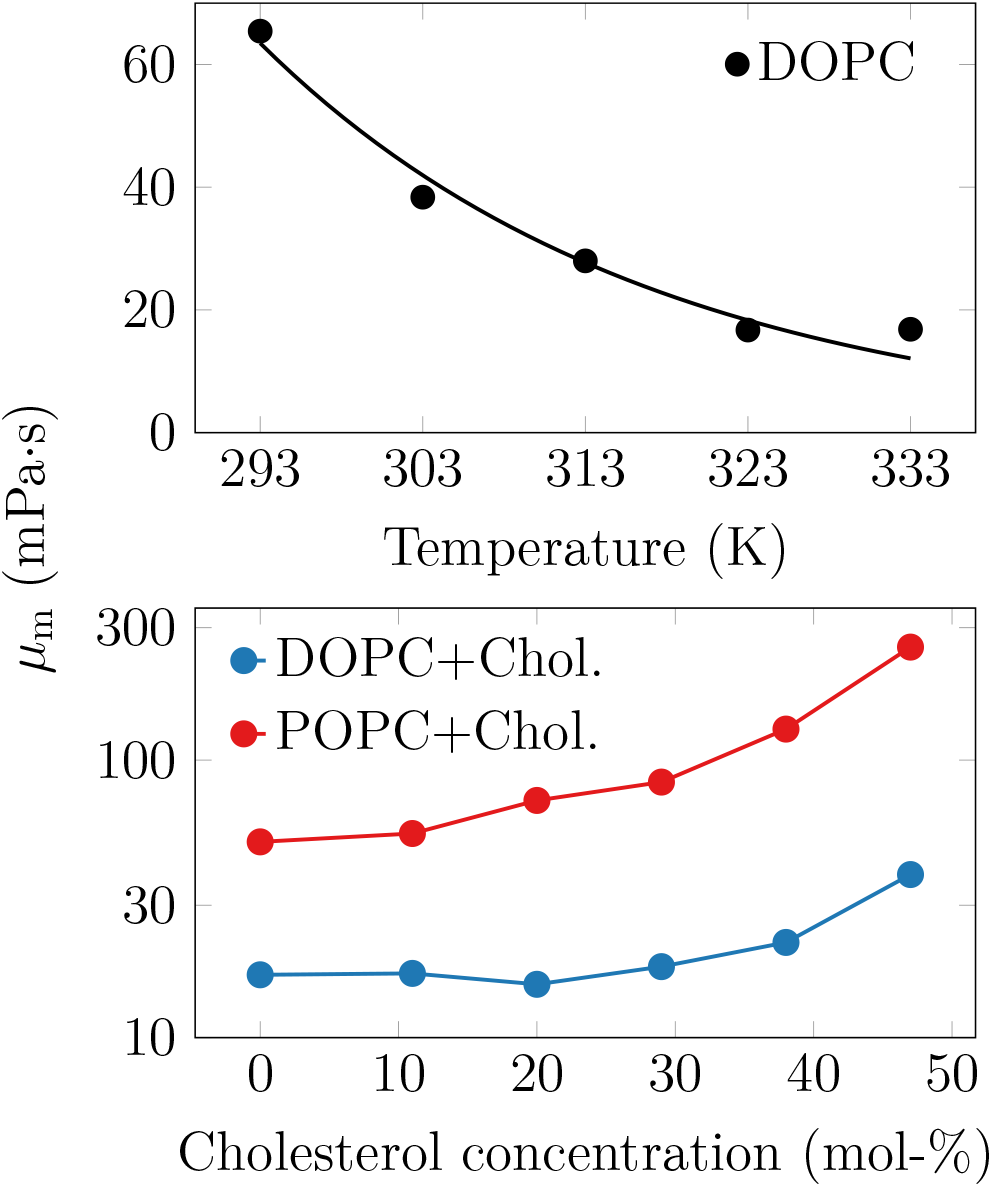
The effect of temperature (top) and cholesterol concentration (bottom) on membrane viscosity, as predicted by atomistic simulations. Note that the DOPC/cholesterol mixtures are simulated at 333 K, and the POPC/cholesterol ones at 298 K, and that in the bottom panel the data are presented on a logarithmic scale.

The experimentally resolved values of membrane shear viscosity for lipid bilayers encompass several orders of magnitude,^22^ and comparing them with the simulation results poses a challenge. Still, at 298 K, the viscosity of DOPC in our simulations is ≈50 mPa s, which is in excellent agreement with the value of 51.3±1.8 mPa s extracted for the same lipid model at 303 K using a non-equilibrium approach.^37^ It also agrees well with the experimental estimates of 49 mPa s obtained using time-resolved fluorescence spectroscopy,^24^ or 41±10 mPa s obtained from the fluorescence of a viscosity-sensitive dye,^71^ both measured at 298 K. For DOPC, Merkel et al. extracted viscosity values of ≈42.9–47.6 mPa s at 318 K, and ≈224 mPa s at 283 K.^72^ Some experiments have reported values that are somewhat larger, like fluorescence lifetime measurements of a rotor that found a value of 228 mPa s,^68^ or an approach based on the simultaneous measurement of rotational and translational diffusion coefficients of lipid-attached ellipsoidal fluorescent microparticles that found a value of ≈500±370 mPa s.^23^ Amador et al. found a value smaller than ≈160 mPa s by shearing a bilayer with optical tweezers, and interpreting the results using numerical simulations.^73^ Higher values of ≈1080±700,^22^ and ≈4400±290 mPa·s^25^ have also been reported for DOPC at room temperature. On the simulation side, somewhat higher values of 65–87 mPa·s are estimated for DOPC at 298 K based on the periodic Saffman–Delbrück model applied to simulations with the same CHARMM36 lipids model.^30^ Note that these membrane shear viscosity values above are either taken from the publications as such, or converted from surface viscosities using membrane thicknesses from our simulations.

The temperature-dependence of viscosity is captured by the activation energy, which is obtained from an Arrhenius analysis of viscosities or diffusion coefficients. Either way, we extracted a value of 29 kJ/mol for our DOPC membranes. We used the diffusion coefficient values extrapolated to infinite size of the simulation box, yet the diffusion coefficients extracted from systems affected by finite-size effects provided similar activation energy estimates. Our result agrees well with the experimental estimates of 27 kJ/mol obtained from NMR diffusion measurements^69^ or 23.5 kJ/mol from fluorescence lifetime measurements,^68^ yet is significantly smaller than another estimate of 54±9 kJ/mol obtained with a viscosity-sensitive probe.^71^

Concluding, these results show that membrane viscosity decreases with increasing temperature as the membrane becomes more fluid. However, in the biologically relevant temperature range, which is usually quite narrow, this change is relatively modest.

### 3.5 The Addition of Cholesterol Significantly Increases Membrane Viscosity

Cholesterol is known to induce membrane ordering and tighter packing of cell membranes, resulting in slower lateral diffusion^74^ (although it should be noted that here we focus on temperatures where the cell membrane remains fluid; at low temperatures, where cholesterol breaks the structure of the gel phase, the situation is just the opposite). While atomistic MD simulation studies have resolved the effect and molecular mechanism of increasing cholesterol concentration on lipid diffusion,^75,76^ the values of diffusion coefficients and their exact trends cannot be compared to experiment. This stems from the fact that increasing cholesterol concentration increases the viscosity of the membrane, as shown by our atomistic simulation data in Fig. 4, and the PBC correction is thus not uniform for such a set of simulations.

The lateral diffusion coefficients for DOPC in a binary mixture with cholesterol, with increasing cholesterol concentration at 333 K, are compared to experimental NMR data^69^ in Fig. S9 in the SI. It is evident that while the results from the largest bilayer simulations agree well with experiment, the PBC-corrected diffusion coefficients are significantly larger than the experimental ones. This already signifies that the viscosity of cholesterol-containing bilayers is too small in simulations. Whereas experimental data display a linear dependence of viscosity on cholesterol concentration, simulation data for DOPC display a kink (see Fig. 4(bottom)), indicating that a phase transition might take place in the system. We also extracted the lateral diffusion coefficients in POPC–cholesterol mixtures at 298 K, and found similar discrepancies between simulation and experiment (see Fig. S9); simulation values were larger when extrapolated to infinite systems at most studied concentrations.

How do our simulation values compare to experimental results? Wu et al. measured a value of 263 mPa s for a 60/40 mixture of DOPC and cholesterol at 298 K, *i.e*. an increase of 16% from the cholesterol-free case.^68^ Chakraborty et al. observed an increase from 4300 to 6800 and to 7800 mPa s upon the addition of 10 or 20 mol% of cholesterol at 298 K, an increase of 59% and 81%, respectively.^25^ Faizi et al. observed an increase from 1080±700 to 1640±1080 mPa·s, *i.e*. an increase of 70% from a pure DOPC bilayer to a 50/50 mixture of DOPC and cholesterol at 298 K.^22^ Compared to these numbers, we observe no effect of cholesterol up to 20 mol%, in disagreement with the results of Chakraborty et al‥^25^ At 40 mol%, we observe an increase of ≈30%, *i.e*. approximately twice the increase found by Wu et al‥^68^ At 50 mol%, our cholesterol-containing bilayer is 130% more viscous than the cholesterol-free one, again overshooting an experimental value of Faizi et al. by a factor of two,^22^ although the error estimates of these experimental values are substantial. While these comparisons are likely affected by the fact that our simulations were performed at a much higher temperature compared to the experiments, even the observed trends differ between our simulations and experiments. If instead of DOPC we consider POPC, whose simulation results correspond to a temperature of 298 K and are thus closer to the experimental conditions, the difference is no longer so noticeable, especially in terms of the trend. These data are discussed below separately.

On the simulation side, Zgorski et al. found that at 323 K, the change from 100 mol% DPPC to a mixture of 55 mol% DPPC, 15 mol% DOPC, and 30 mol% cholesterol essentially doubled the viscosity.^37^ However, comparison to our DOPC/cholesterol and POPC/cholesterol mixtures is not straightforward, since the interactions of cholesterol with the saturated chains of DPPC lead to a larger condensation effect than with POPC or DOPC, and thus likely has a larger impact on viscosity.^77^

Unfortunately, no experimental data exists for the mixture of POPC and cholesterol. Still, we obtained very similar viscosity values for DOPC (51.9 mPa s) and POPC (50.7 mPa s, interpolated estimate) at 298 K, respectively. However, experimentally, POPC viscosity was found to be more than twice larger than that of DOPC at this temperature.^22^ Due to the lower temperature (298 K vs. 333 K), our POPC–cholesterol membranes are more viscous than the DOPC–cholesterol ones. At the largest studied cholesterol concentration, the POPC–cholesterol mixture reaches a viscosity value of 256 mPa·s, some 5-fold higher than the cholesterol-free case. In experiments for the DOPC–cholesterol mixture at 298 K, a similar amount of cholesterol only resulted in an increase of viscosity by 16%,^68^ suggesting that the effects of cholesterol are greatly exaggerated in the CHARMM36 force field.

Concluding, cholesterol weakens the fluidity of cell membranes and increases their viscosity, but the details of this feature clearly depend on the thermodynamic conditions.

### 3.6 Acyl Chain Length and Unsaturation Level Dictate Membrane Viscosity

Next, we shed light on the effects of acyl chain length and saturation level. These trends are shown in Fig. 5. An increase in chain length or saturation level increases viscosity, as expected. For the lipids considered here, each methyl group increases viscosity on average by 3.4±1.9 mPa·s. Similarly, each double bond decreases viscosity by 1.9±0.7 mPa·s. In the analysis regarding the contribution of double bonds, data for DOPC (18:1,18:1) were left out as they were outliers in Fig. 5.

**Figure 5:**
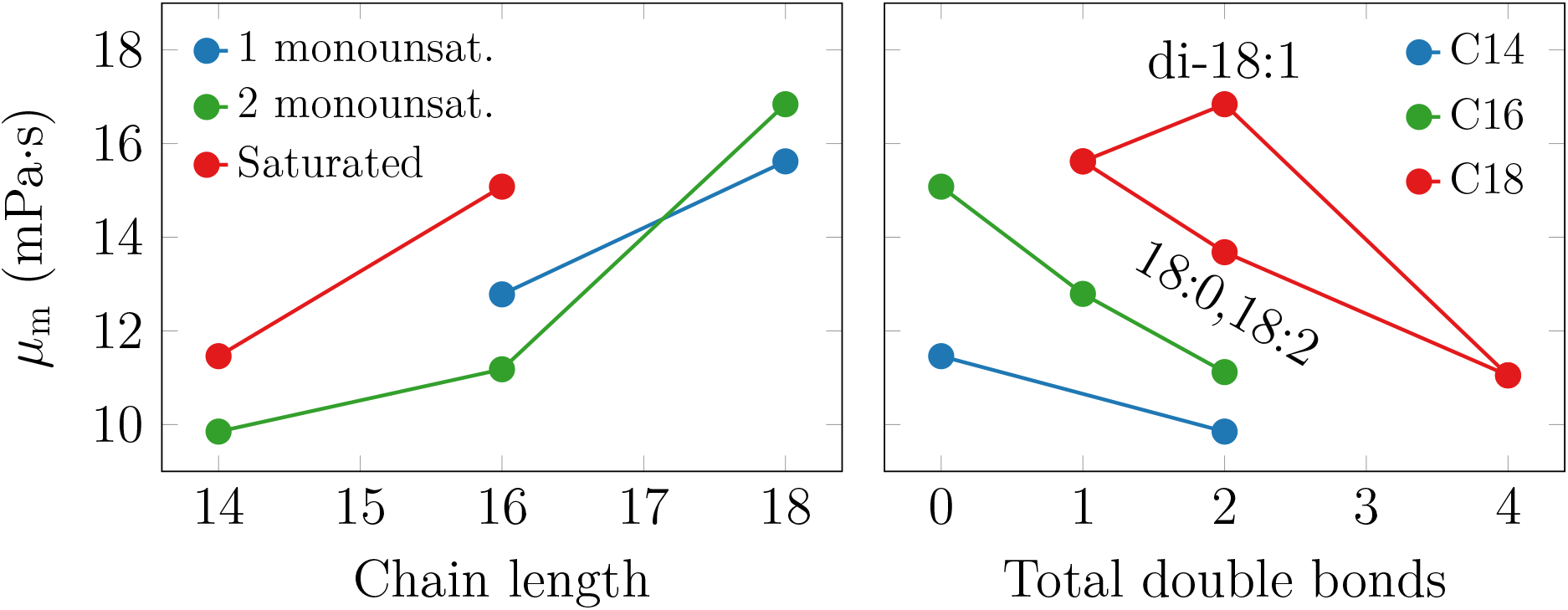
Effects of chain length (left) and chain unsaturation (right) on membrane viscosity. The trends are fairly evident, yet DOPC (18:1,18:1) deviates from the behavior of the other systems.

Recent fluorescence microscopy experiments on ellipsoidal microparticles also considered the effect of chain length for a series of PC lipids ranging from 14 to 20 carbons.^23^ In these measurements, the two chains were of equal length, and one of them contained a double bond. Interestingly, Jahl and Parthasarathy found a very small length-dependence of surface viscosity, yet the membrane shear viscosities actually demonstrated no dependence on acyl chain length, in contrast to our findings. Here, to convert the surface viscosity values to shear viscosity values, we assumed, based on the thicknesses of simulated PYPC (16:0,16:1) and SOPC (18:0,18:1) membranes, that the membrane becomes 0.36 nm thicker with the addition of two methyl groups in each acyl chain.

There are no systematic studies on the effect of chain unsaturation on membrane viscosity. Zgorski et al. extracted values for DOPC and DPPC from simulations of the same lipid model as we use here, albeit not at the same temperature. They obtained 31.9±1.3 mPa s for DPPC at 323 K, which is quite a lot smaller than the value of 51.3±1.8 mPa s extracted for DOPC at 303 K.^37^ On the experimental side, Faizi et al. measured the viscosities of SOPC and DOPC bilayers at 298 K, and observed a decrease of 60 % when the second double bond was introduced.^22^ Curiously, our simulations actually predicted very similar values to SOPC and DOPC. Nojima and Iwata measured viscosities for DMPC to be ≈2-fold more viscous than DOPC, whereas our simulations found DOPC to be more viscous by ≈50%. Finally, Merkel et al. reported very similar viscosity values for DMPC and DOPC at 318 K.^72^ These numbers might again depend on temperature, but performing a temperature scan with all the considered compositions would be simply unfeasible.

We conclude that based on our simulation results, changes in the unsaturation levels of lipid hydrocarbon chains do not significantly change the viscosity of cell membranes.

## 4 Conclusions

Molecular dynamics simulations have the ability to provide insight into how interactions among membrane constituents affect their dynamics. They can explain trends observed in experiments that probe diffusion or related phenomena, such as the rate of diffusion-limited reactions. However, the use of periodic boundary conditions prevents a direct comparison of lateral or rotational diffusion coefficient values with those measured by experiment. Instead, these values need to be first extrapolated to an infinitely large system, corresponding to the experimental setup. Since temperature, cholesterol concentration, or lipid acyl chain length all affect membrane viscosity, the size of the related PBC-corrections vary significantly between different systems. Thus, even the trends observed for a set of systems simulated with PBCs cannot be directly and reliably compared to experiment. The PBC-corrections are easily applied if the membrane viscosity is known. However, this is often not the case as very few such values have been reported by simulation studies.

Here, we have extracted membrane viscosities using several different approaches based on the lateral (translational) and rotational diffusion of membrane proteins and lipids in equilibrium simulations. We first applied multiple approaches to coarse-grained simulations of protein-containing membranes, and found the extracted values to be relatively consistent. Importantly, the PBC-corrections provide a straightforward way of extracting the membrane viscosity values, albeit they require performing simulations of the same membrane composition in systems of multiple sizes. Then, using the most suitable approaches, we demonstrated that for a single protein in a bilayer, the viscosity felt by lateral and rotational diffusive motions is largely similar. This suggests that the contrasting results found in studies of cells might be caused by, *e.g*., protein oligomerization or perturbations due to actin cytoskeleton. We also extracted viscosity values for a protein-crowded lipid membrane, and found it to be significantly more viscous than a protein-dilute membrane, yet in a strongly temperature-dependent manner. In all cases, activation energies extracted from lateral and rotational diffusion coefficients as well as from membrane shear viscosities were similar, indicating that the explicit temperature dependencies of the SD models are dominated by the implicit exponential temperature dependence of the membrane shear viscosity.

We then systematically extracted shear viscosity values for phosphatidylcholine lipids that differed in their acyl chain length and unsaturation level from all-atom simulations. Additionally, we studied the effect of temperature and cholesterol content on membrane viscosity. While the trends observed in simulation data were fairly clear, the comparison to experiment revealed severe shortcomings of the simulation model. These findings highlight how important it is to consider dynamics in the parameterization of lipid models. Popular lipid models like CHARMM36 capture the structural effects brought about by the changes in cholesterol concentration,^54^ temperature,^78,79^ or the properties of the acyl chains.^79^ However, the experimental values or even the trends in membrane shear viscosity are not captured by CHARMM36. This further indicates that even if diffusion coefficients from CHARMM36 simulations are corrected for PBC-induced effects, their comparison to experiment should still be performed with care. Finally, we hope that our study serves as a summary of methods that are available for extracting membrane viscosity, and also as a reference for viscosity values for different lipid membranes in simulations.

## Acknowledgement

We thank CSC–IT Center for Science for computational resources. MJ thanks the Academy of Finland (Postdoctoral grant no. 338160) and the Emil Aaltonen foundation for funding. IV thanks the Helsinki Institute of Life Science (HiLIFE) Fellow program, Human Frontier Science Program (HFSP, project no. RGP0059/2019), Sigrid Jusélius Foundation, and Academy of Finland (project ID: 335527, 331349) for financial support. We thank Victor Cruces Chamorro for calculating water viscosity with the Green–Kubo approach.

