## Supplementary Information for "Protein Crowding and Cholesterol Increase Cell Membrane Viscosity in a Temperature Dependent Manner"

### Supporting Information for: Protein Crowding and Cholesterol Increase Cell Membrane Viscosity in a Temperature Dependent Manner

Balázs Fábián,[*,†,‡] Ilpo Vattulainen,[¶] and Matti Javanainen[*,§,†,||]

†*Institute of Organic Chemistry and Biochemistry of the Czech Academy of Sciences, Flemingovo nám. 542/2, CZ-16000 Prague 6, Czech Republic*

‡*Current address: Department of Theoretical Biophysics, Max Planck Institute of Biophysics, Max-von-Laue Straße 3, 60438, Frankfurt am Main, Germany*

¶*Department of Physics, University of Helsinki, FI-00014 University of Helsinki, Helsinki, Finland*

§*Institute of Biotechnology, University of Helsinki, FI-00014 University of Helsinki, Helsinki, Finland*

||*Computational Physics Laboratory, Tampere University, Tampere, Finland*

### 1 Supplementary Methods

#### 1.1 All-Atom Simulations

We used the CHARMM36 lipid model for both phospholipids[1] and cholesterol.[2] The cholesterol-free membranes contained either 64, 256, or 1024 phospholipids, whereas in cholesterolcontaining membranes, the cholesterol molecules were added without removing any phospholipids. The membranes were solvated (50 waters per lipid) by the CHARMMM-specific TIP3P model.[3,4]

We used the simulation parameters recommended for CHARMM36 in GROMACS.[5] These simulation parameters, as well as the topologies, were obtained in GROMACS formats from CHARMM-GUI.[5] A time step of 2 fs was enabled by constraining bonds involving hydrogens by P-LINCS.[6,7] Calculation of non-bonded interactions was facilitated by buffered Verlet lists that tracked atomic neighbors.[8] For electrostatics, we used the smooth PME approach to include the long-range contributions.[9,10] Van der Waals and steric interactions were modeled by the Lennard-Jones potential with a cut-off of 1.2 nm, while the forces were switched to 0 between 1.0 and 1.2 nm to avoid discontinuity at the cut-off distance. The temperatures of the membrane and the solvent were separately coupled to the Nosé–Hoover thermostat[11,12] with a time constant of 1 ps. The target temperatures varied, as explained in the main text. The Parrinello–Rahman barostat[13] with a time constant of 5 ps, a compressibility of $4.5\times10^{-5}$ bar$^{-1}$, and a reference pressure of 1 bar were used to maintain a constant pressure both in the plane of the membrane, as well as normal to it. All simulations were performed with the GROMACS 2020 package.[14,15]

### 2 Supplementary Results

#### 2.1 Results From Coarse-Grained Simulations

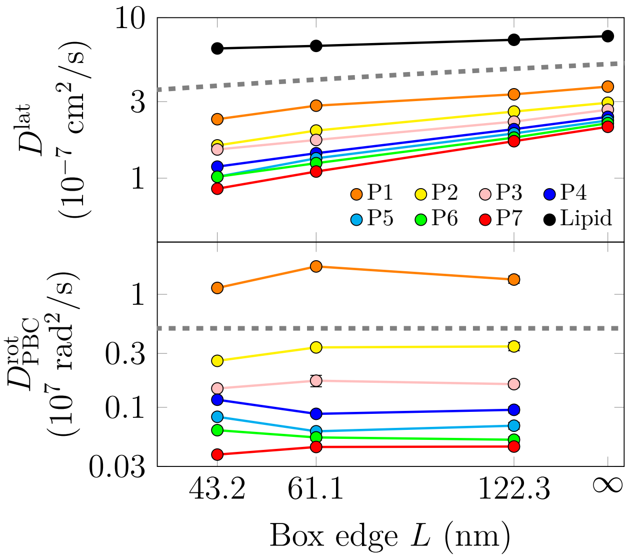


Figure S1: Dependence of lateral (top) and rotational (bottom) diffusion coefficients on membrane size extracted from the CG simulations where all proteins are present at once (Systems "P-" in Table 1, *i.e.* Set CG-1). For the former, a logarithmic $D^{\mathrm{lat}} \sim \ln(L)$ dependence is found (dashed line),[16] whereas for the latter, no size dependence is observed (dashed line), in line with Ref. 17. All shown figures for rotational diffusion are affected by PBCs, albeit the effect is minor for systems of this size.[17] For lateral diffusion, in addition to $D^{\mathrm{lat}}_{\mathrm{PBC}}$, we also plot the values at $L = \infty$ ($D^{\mathrm{lat}}_{\infty}$), extrapolated from Eq. 3 in the main text. Importantly, the scaling relation also holds well for a lipid molecule, which experiences a very different environment due to inter-leaflet coupling. This independence of the correction on the type of studied molecule undergoing diffusive motion is a convenient feature of the hydrodynamic correction.[18]

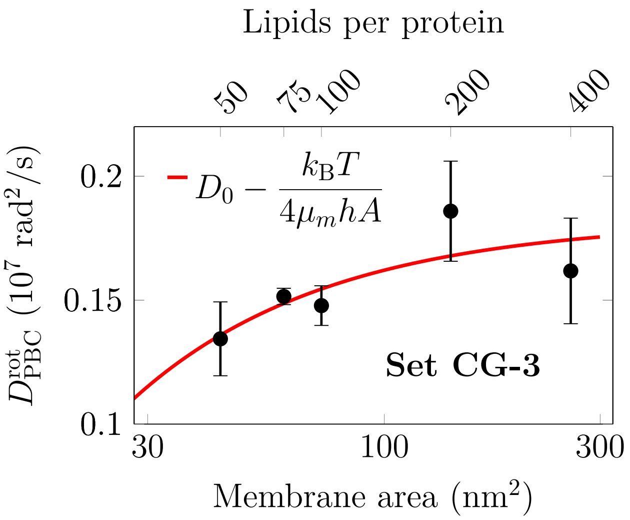


Figure S2: Dependence of rotational diffusion coefficients on membrane size in the CG simulations. Here, the medium-sized protein "P3" with an effective radius of 2.44 nm was used in a single-protein setting ("S-3-" in Table 1). The size dependence is satisfactorily captured by Eq. (4), and is hence in line with Ref. 17.

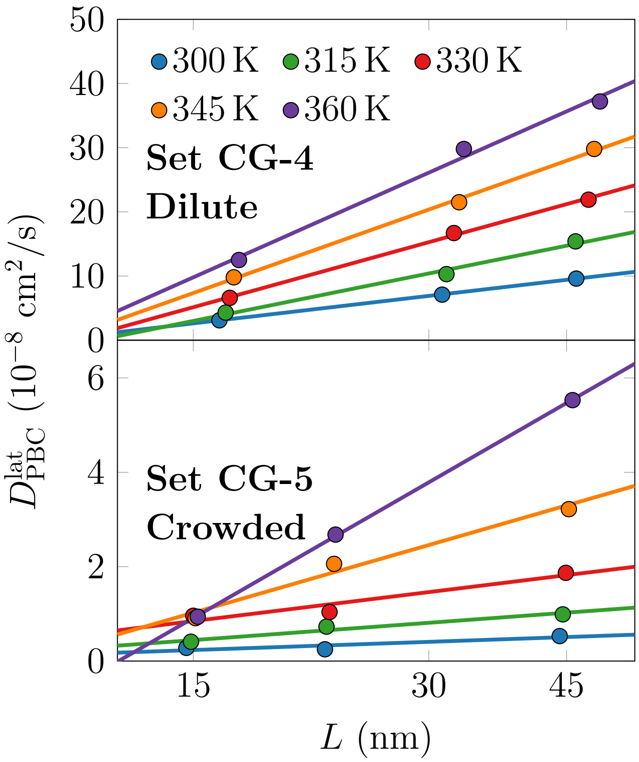


Figure S3: The dependence of lateral diffusion coefficients of proteins on the system size in dilute and crowded CG systems. The top panel shows data for dilute single-protein systems ("S-$4^{400}$", "S-$4^{900}$,", and "S-$4^{3600}$", *i.e.* Set CG-4 in Table 1 in the main text), whereas the bottom panel shows data for crowded multi-protein systems ("M-$4^{400}$, M-$4^{900}$, and M-$4^{3600}$", *i.e.* Set CG-5 in Table 1 in the main text). The lines show fits of logarithmic dependence on the box size ($D^{\mathrm{lat}} \sim \ln L$). Note the logarithmic scale on the abscissa.

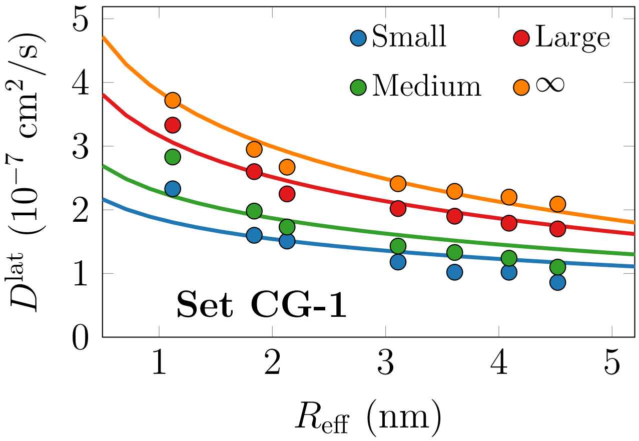


Figure S4: The dependence of lateral diffusion coefficients ($D^{\rm lat}_{\rm PBC}$) on the effective protein radius in the "P-" systems of three different sizes (Set CG-1), as well as in the infinite system size ($D^{\rm lat}_{\infty}$) extrapolated using Eq. (3) in the main text. These values are also shown in Fig. S1 as a function of system size. The solid lines show fits of the SD model, Eq. (1), to the data. Not surprisingly, only the diffusion coefficients of the infinite system size follow the SD-like $D^{\rm lat} \propto \ln(R)$ dependence.

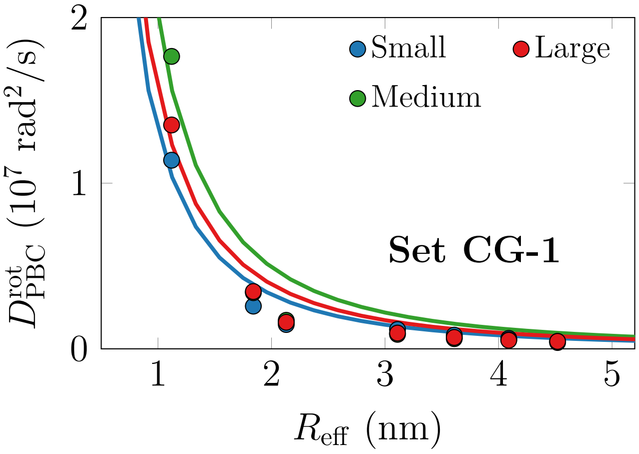


Figure S5: The dependence of rotational diffusion coefficients on the effective protein radius in the "P-" systems of three different sizes (Set CG-1). With these system sizes, the PBC-correction for rotational diffusion is minuscule (see also the bottom panel of Fig. S1). The solid lines show fits of the SD model for rotational diffusion, Eq. (2), to the simulation data.

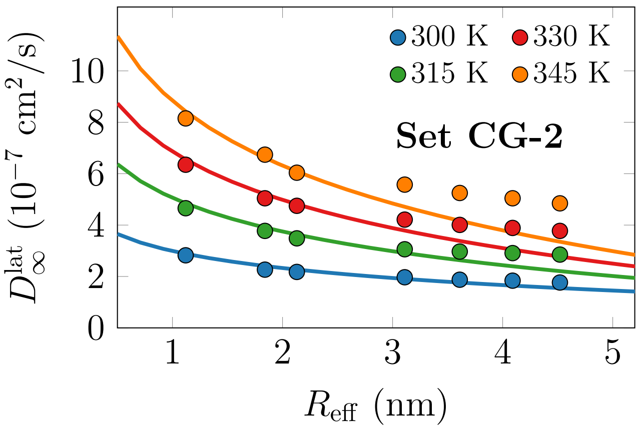


Figure S6: The dependence of lateral diffusion coefficients on the effective protein radius in the S-3 systems (of Set CG-2) at four different temperatures. All the shown values have been corrected for finite-size effects using the viscosity extracted from fits to finite-size dependency of lateral diffusion (Set CG-4 & Figs. S1 and 2) using Eq. (3). The solid lines show fits of the SD model for lateral diffusion, Eq. (1), to the data. The data measured at lower temperatures seem to better agree with the fits.

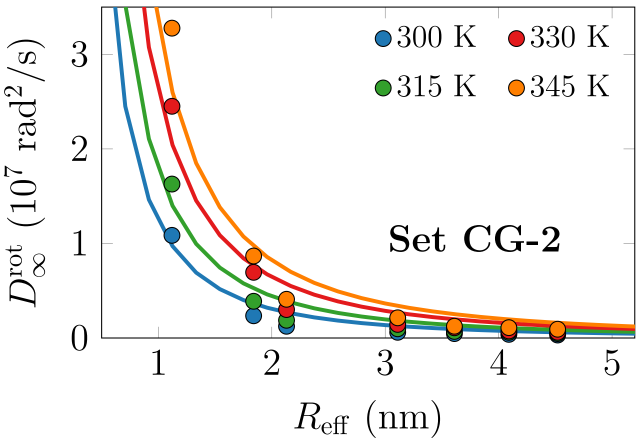


Figure S7: The dependence of rotational diffusion coefficients on the effective protein radius in the S-3 systems (of Set CG-2) at four different temperatures. All the shown values have been corrected for finite-size effects using the geometric correction factor of Eq. (4). The solid lines show fits of the SD model for rotational diffusion, Eq. (2), to the data.

#### 2.2 Results From All-Atom Simulations

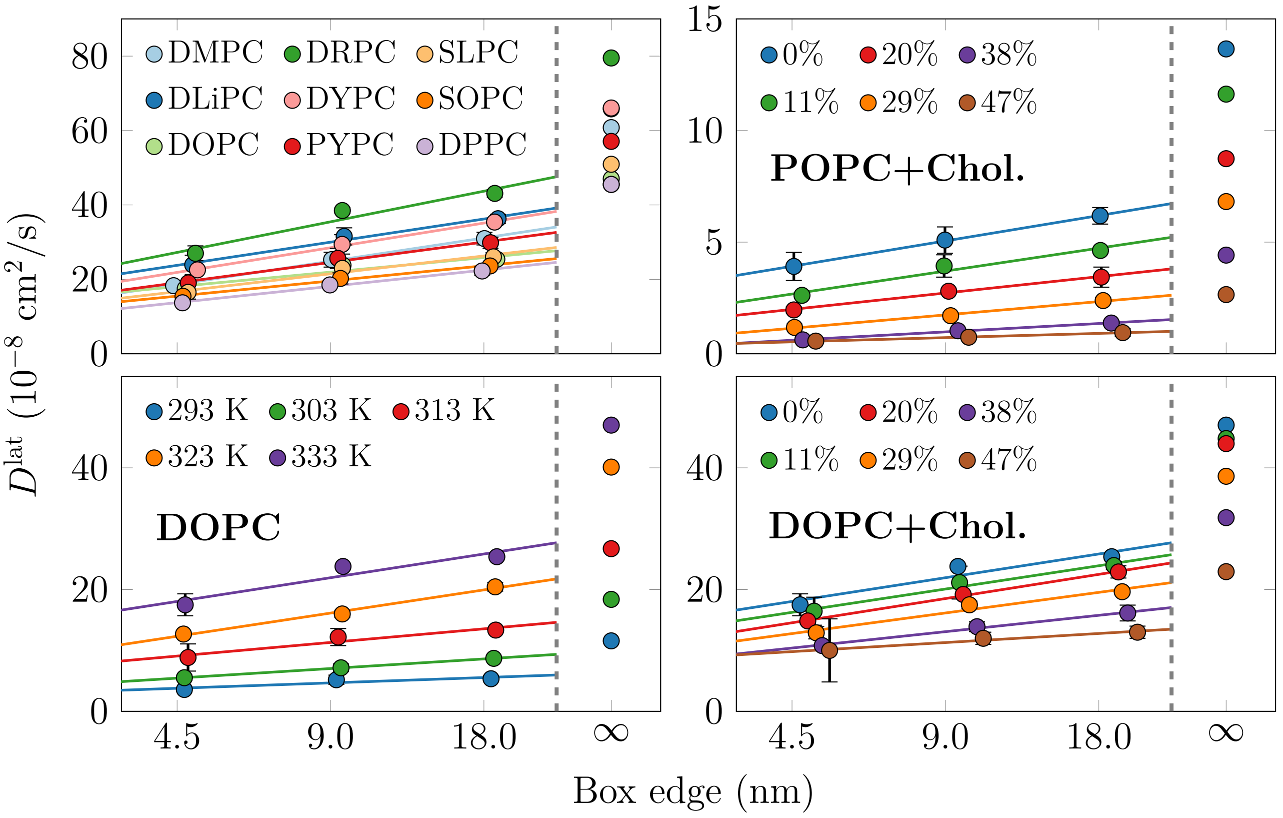


Figure S8: Lateral diffusion coefficients ($D^{\text{lat}}_{\text{PBC}}$) of lipids in all-atom simulations as a function of lateral size of the simulation box. The three markers show the values calculated from simulations performed using three different system sizes. The solid lines show fits of Eq. (3) to these three data points, and the marker at $\infty$ shows the value extrapolated to infinite box size from the fits ($D^{\text{lat}}_{\infty}$). Data are shown for phosphatidylcholines with different acyl chains (top left panel) at 333 K, for DOPC at different temperatures (bottom left panel), for POPC mixed with cholesterol at 298 K, and for DOPC mixed with cholesterol at 333 K.

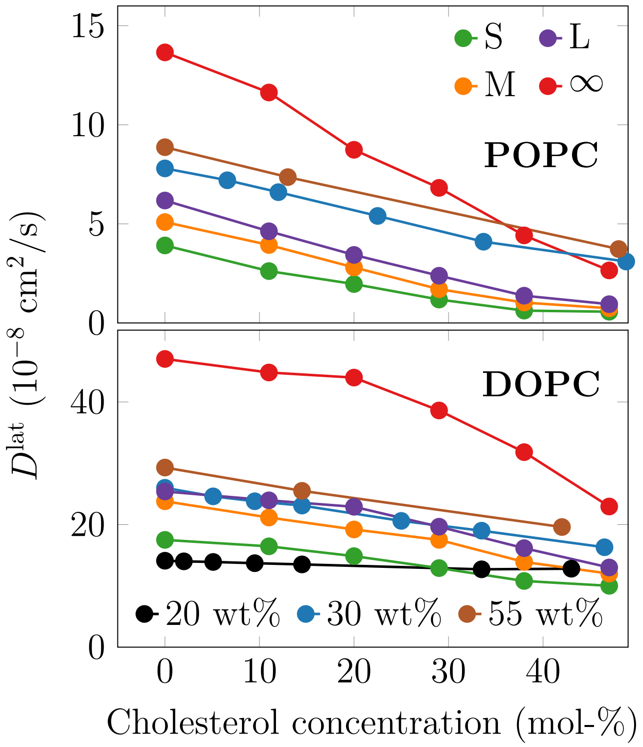


Figure S9: Lateral diffusion coefficients ($D^{\mathrm{lat}}_{\mathrm{PBC}}$) of POPC (298 K, top) and DOPC (333 K, bottom) as a function of cholesterol concentration, based on atomistic MD simulations. Values extracted from the simulation systems of three different sizes are shown; small (S), medium (M), and large (L). Moreover, values extrapolated to infinite ($D^{\mathrm{lat}}_{\infty}$) system sizes using Eq. (3) are shown. Experimental data have been measured using the pulsed field gradient NMR approach at multiple hydration levels (20, 30, and 55 weight% of water in the sample).[19,20] These values typically converge at the largest shown hydration level of 55 weight%.[20]

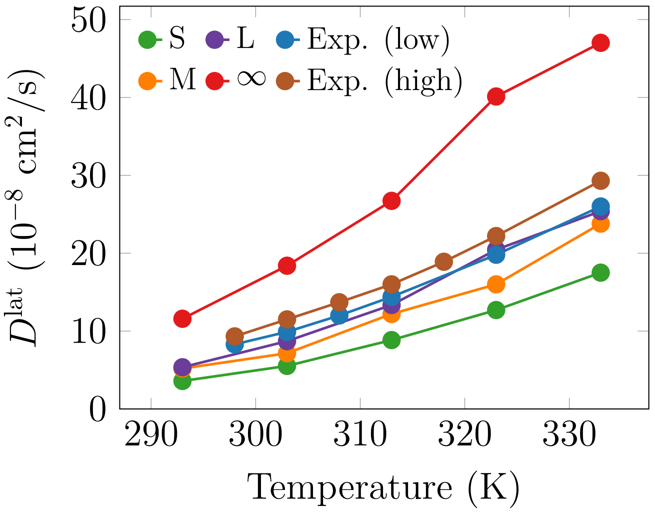


Figure S10: Lateral diffusion coefficients of DOPC ($D^{\mathrm{lat}}_{\mathrm{PBC}}$) as a function of temperature extracted from the atomistic simulation data of three different system sizes (small "S", medium "M", and large "L"), as well as from the data extrapolated to infinite system size ($D^{\mathrm{lat}}_{\infty}$) using Eq. (3) (∞).

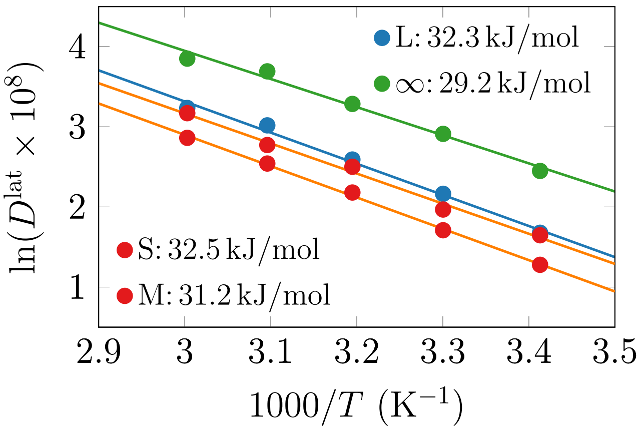


Figure S11: Lateral diffusion coefficients of DOPC shown in the Arrhenius description (Eq. (7)), *i.e.* the logarithm of the (unitless) diffusion coefficient is shown as a function of inverse temperature. Atomistic simulation sata are shown for three different system sizes ($D^{\mathrm{lat}}_{\mathrm{PBC}}$, small "S", medium "M", and large "L"), as well as for the extrapolated infinite system $D^{\mathrm{lat}}_{\infty}$) size using Eq. (3) (∞). The activation energies extracted using the Arrhenius form are also shown in the legend labels.
